# Primary Human Hepatocytes Maintain Long-term Functions in Porous Silk Scaffolds Containing Extracellular Matrix Proteins

**DOI:** 10.1101/2020.01.30.927814

**Authors:** David A. Kukla, Whitney L. Stoppel, David L. Kaplan, Salman R. Khetani

**Affiliations:** Department of Bioengineering, University of Illinois at Chicago, Chicago, IL; Department of Biomedical Engineering, Tufts University, Medford, MA

**Keywords:** 3T3-J2 fibroblasts, Matrigel, type I collagen, porous scaffolds, silk fibroin

## Abstract

The shortage of donor organs for transplantation has prompted the development of alternative implantable human liver tissues; however, the need for a clinically viable liver tissue that can be fabricated using physiologically-relevant primary human hepatocytes (PHHs) is unmet. Purified silk proteins provide desirable features for generating implantable tissues, such as sustainable sourcing from insects/arachnids, biocompatibility, tunable mechanical properties and degradation rates, and low immunogenicity upon implantation; however, the utility of such scaffolds to generate human liver tissues using PHHs remains unclear. Here, we show that the incorporation of type I collagen during the fabrication and/or autoclaving of silk scaffolds was necessary to enable robust PHH attachment/function. Scaffolds with small pores (73 +/- 25 µm) promoted higher PHH functions than large pores (235 +/- 84 µm). Further incorporation of growth-arrested 3T3-J2 fibroblasts into scaffolds enhanced PHH functions up to 5-fold for 5 months in culture, an unprecedented longevity, and functions were better retained than 2D configurations. Lastly, encapsulating PHHs within Matrigel™ while housed in the silk/collagen scaffold led to higher functions than Matrigel or silk/collagen alone. In conclusion, porous silk scaffolds are useful for generating long-term PHH +/- fibroblast tissues which may ultimately find applications in regenerative medicine and drug development.

## INTRODUCTION

The diverse functions of the liver can be severely compromised by several diseases. In particular, drug-induced liver injury (DILI) accounts for ∼50% of acute liver failures and is the leading cause of drug attrition [1], while hepatitis B and C viruses chronically infect ∼400 million and ∼170 million people, respectively, and can lead to fibrosis, cirrhosis, and hepatocellular carcinoma (HCC), the third leading cause of deaths from cancer globally [2]. Additionally, non-alcoholic fatty liver disease is on an epidemic rise and can also lead to fibrosis, cirrhosis, and HCC [2]. Overall, chronic liver disease affects >500 million people, causing 2% of all deaths globally [3, 4]. Currently, orthotopic liver transplantation is the only treatment for liver failure shown to directly alter mortality [5]. However, there is a severe shortage of viable donor livers to satisfy increasing demand; for example, 8,250 liver transplants were performed in the US in 2018 whereas there are almost 14,000 patients currently on the waiting list (American Liver Foundation). To address these shortages and enable new therapeutic advances, there is a need for the development of alternative cell-based therapies that can serve as a bridge to liver transplantation and eventually as a complete organ replacement. Additional understanding of the key cell types and culture parameters that enable long-term *in vitro* function of engineered liver tissue is critical for the achievement of new therapeutic advances.

Given significant species-specific differences in liver functions (e.g. drug metabolism pathways) and drug toxicity outcomes, human liver cells are now being utilized for fabricating platforms useful for cell-based therapies and/or drug screening [6]. However, immortalized cell lines display abnormal growth and low liver functions such as drug metabolism capacity [7], while protocols to generate autogenic induced pluripotent stem cell-derived hepatocyte-like cells require further improvements to enable adult-like functions [8]. Therefore, primary human hepatocytes (PHHs), that perform the majority of functions in the liver *in vivo*, are considered the gold standard for building human liver platforms. While PHHs are in limited supply, efforts are underway to harness their tremendous *in vivo*-like growth potential using small molecules and/or growth factors *in vitro* [9] or expansion *in vivo* in rodents stimulated to produce regenerative factors [10]. Since infusion of isolated PHHs into patients has not shown long-term clinical benefit due to limited cell engraftment and immune-mediated reaction, a biomaterial scaffold housing viable PHHs long-term either in extracorporeal devices or for implantation with immunomodulatory properties will be necessary for sustainable therapy [5].

Others have previously fabricated 3-dimensional (3D) implantable liver constructs containing PHHs housed within different biomaterials. In particular, Kostadinova et al seeded liver stromal cells and PHHs onto nylon scaffolds and measured 77 days of liver function [11]; however, while useful for drug screening, the non-biodegradable nylon is not suitable for implantation. In contrast, Organovo, Inc. bio-printed a human liver-like tissue containing adjacent compartments housing PHHs and liver stromal cells encapsulated within a degradable bioink that displayed 28 days of function [12]; however, the long-term (months) functionality of such tissues is unclear and the use of such mechanically weak and rapidly degradable tissues for implantation may not be desirable. Similarly, PHHs encapsulated into alginate microbeads functioned for 3 days *in vitro* and improved liver functions in rodents with acute liver failure for ∼7 days [13]; such short-term outcomes are likely due to the bioinert alginate not being suitable for long-term culture without further cross-linking [14], which can impact cell signaling and/or cell viability [15, 16]. In contrast to naturally-derived alginate, Chen et al used synthetic and non-biodegradable polyethylene glycol diacrylate (PEGDA) to encapsulate PHHs and supportive fibroblasts via UV crosslinking, though induction of high PHH functions for 8 days required PEGDA modification with RGDS [17]. Furthermore, intraperitoneal implantation of the PEGDA-RGDS constructs into rodent models showed retention of PHHs and penetration of host vasculature after several weeks [17]. However, it is not clear if a) PHHs can maintain high levels of functions within PEGDA-RGDS scaffolds for longer than 1-2 weeks *in vitro* towards providing a longer shelf-life for on-demand implantation into recipients and b) UV cross-linking that is necessary to encapsulate cells into PEGDA induces any long-term genotoxicity and/or other signaling changes in PHHs. Lastly, besides being mechanically weak, PEGDA requires modification with protease-activated peptides to render it biodegradable for integration with host tissue, which adds to fabrication complexity and cost. Therefore, while the studies above have demonstrated proof-of-concept for generating PHH-laden biomaterial scaffolds for implantation purposes, there is a need to further explore alternative biomaterials that can be used to fabricate optimal cell-based therapies.

Scaffolds made of silk fibroin protein are known to be biocompatible for long-term (months) culture of cells *in vitro* without causing cell-specific signaling, without needing any chemical or UV-based crosslinking, and without degrading/collapsing in cell culture medium or under perfusion within bioreactors; additionally, silk scaffolds can be autoclaved for contaminant-free culture [18, 19]. From a manufacturing perspective, silk scaffolds can be a) fabricated using proteins secreted by many insect/arachnid species (e.g. silkworms and spiders), b) molded into a wide range of forms (e.g. hydrogels and sponges), c) tuned for porosity while maintaining robust bulk mechanical properties and *in vivo* degradation rates, and d) combined with other materials (e.g. ceramics and extracellular matrix or ECM proteins) to create composites that modulate cell responses [20]. When implanted *in vivo*, silk sponges and other formats can induce a transient (weeks) mild and beneficial inflammatory response that induces vascular ingrowth but does not activate the adaptive immune response nor the formation of a permanent fibrotic capsule [21]. Furthermore, silk degradation into non-toxic byproducts *in vivo* can be extended for months to allow native tissue remodeling. While silk scaffolds, in some cases with incorporated ECM proteins/gels, have been used to generate long-term tissue constructs for several different tissue types [20, 22] as well as for the culture of rat hepatocytes +/- hepatic stellate cells for 2-3 weeks [23-26], it is not clear if silk scaffolds can support long-term PHH functions. Therefore, here we sought to determine for the first time the long-term function of PHHs in porous silk scaffolds +/- type I collagen. We examined the role of collagen I and Matrigel™ in improving cell adhesion and retention within the sponges; we further explored how the inclusion of supportive fibroblasts in silk scaffolds with an entrapped ECM gel affects PHH functions for up to 5 months *in vitro*.

## METHODS

### Preparation of porous silk sponges (scaffolds)

Silk fibroin solution was prepared as previously reported [18, 27, 28]. Briefly, pure silk fibroin was extracted from *Bombyx mori* cocoons by degumming 5 grams of fibers in sodium carbonate solution (0.02M) to remove sericin. Degummed fibroin fibers were rinsed in distilled water and solubilized in aqueous lithium bromide (9.3M) for 4 hours at 60°C. The solution was dialyzed against deionized water until the conductivity of the dialysis water was <10 μS cm^-1^. The solubilized silk solution was centrifuged to remove insoluble particulates. The final concentration was 3% wt/v silk solution. To fabricate collagen Incorporated silk, 1 mg/mL rat tail collagen I (Corning Life Sciences, Tewksbury, MA) in distilled water was added to 3% wt/v silk fibroin solution. Aqueous silk solutions +/- incorporated collagen were dispensed into wells of a 6-well plate. Silk +/- incorporated collagen was frozen overnight at −20°C and lyophilized at either − 20°C or −80°C to create a porous network with either large (235 µm +/- standard deviation of 84 µm) or small (∼73 µm +/- standard deviation of 25 µm) pores, respectively. Dry scaffolds were removed and rendered insoluble by autoclaving at 121°C for 20 minutes at 15 psi in either 1X phosphate-buffered saline (PBS) or 1 mg/mL collagen in 1X PBS to induce β-sheet formation. Scaffolds were cut to size using 6 mm biopsy punches (h x w: 4.4 +/- 0.3 mm x 5.7 × 0.3 mm) and stored in 1X PBS at 4°C until use for cell culture. Silk scaffolds without any collagen incorporation via the processing steps above (i.e. scaffolds autoclaved in 1X PBS alone) were placed in 1 mg/mL collagen in 1X PBS for 2 hours at 37°C to enable passive collagen adsorption.

### Cell culture

Cryopreserved PHHs were purchased from vendors permitted to sell products derived from human organs procured in the United States of America by federally designated Organ Procurement Organizations (BioIVT, Baltimore, MD, and Lonza, Walkersville, MD). PHH lots included HUM4055A (54-year-old female Caucasian) from Lonza and EJW (29-year old female Caucasian) from BioIVT. PHHs were thawed, counted, and viability was assessed as previously described [29]. 3T3-J2 murine embryonic fibroblasts were passaged as previously described and for some experiments, growth arrested by incubating in culture medium containing 1 μg/mL mitomycin-C (Sigma-Aldrich) for 4 hours before trypsinization.

Prior to cell seeding, 48-well culture plates were coated with 5% w/v Pluronic™ F-127 (Sigma-Aldrich, St. Louis, MO) to make the surface of the tissue culture plastic non-adherent to cells [30]. Furthermore, all types of porous silk scaffolds described above were placed in hepatocyte culture medium containing 10% fetal bovine serum (FBS, Thermo Fisher, Waltham, MA) overnight to promote protein binding to the silk surface before cell seeding. Other components of the hepatocyte culture medium were described previously [29].

Cell suspensions were prepared in culture medium containing either PHH only (4e6 cells/mL), co-cultures of PHHs and growth-competent 3T3-J2 fibroblasts, or co-cultures of PHHs with growth-arrested 3T3-J2 fibroblasts; the ratio between the two cell types was kept at 1:1 (8e6 total cells/mL). For certain conditions, cells were suspended in either 4 mg/mL rat tail collagen I or growth factor reduced Matrigel™ (Corning Life Sciences) solutions. Silk scaffolds were partially dehydrated by manually squeezing the hepatocyte culture medium out and then the scaffolds were placed within the Pluronic F-127-coated wells of the 48-well plate. The above-mentioned cell suspensions were seeded onto the partially dehydrated silk scaffolds at 125 µL per scaffold (500K PHHs +/- 500K 3T3-J2 fibroblasts); the dehydrated silk scaffolds rapidly absorbed the cell suspensions (**Fig. 1**). Cells were then allowed to attach to the scaffold for ∼3-4 hours at 37°C; cell suspension that pooled to the bottom of the well was periodically pipetted back onto the scaffolds during this timeframe to promote further cell attachment to the scaffold. After cell attachment, additional culture media (500 µL/well) was added to each well. For control conditions, cell suspensions in the collagen or Matrigel solutions were pipetted directly into the wells and allowed to polymerize as hydrogels in the absence of silk at 37°C. All cultures were maintained *in vitro* for up to ∼5 months with culture medium changes every 4 days.

**Figure 1.**
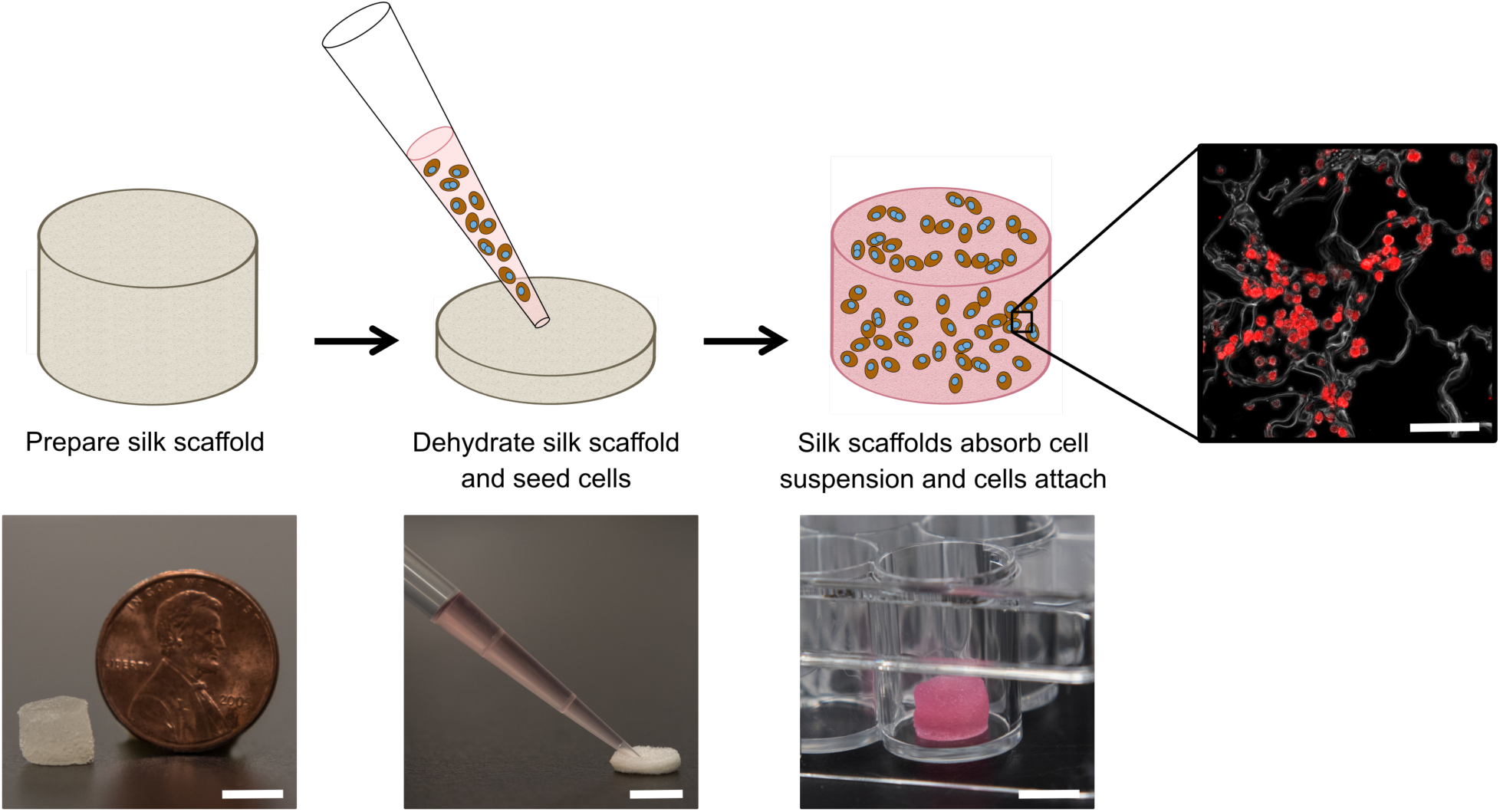
Culture of cells within porous silk sponges (scaffolds). Left to right: Insoluble silk scaffolds containing type I collagen (from rat tails) were fabricated and autoclaved per the protocols described in methods and then partially dehydrated by manually squeezing the hepatocyte culture medium out of the pores. The scaffolds were subsequently placed within the wells of a 48-well plate which was pre-coated with Pluronic F-127 to prevent cell attachment to the plastic. A cell suspension of PHHs +/- 3T3-J2 fibroblasts was seeded onto the partially dehydrated silk scaffolds, which wicked up the cell suspension and allowed cells to attach over 3-4 hours at 37°C to the collagen within the silk scaffolds. Once cells attached, additional cell culture medium was added to the wells and replaced every 4 days with fresh medium. Culture medium was collected for supernatant-based assays and the cell-laden scaffolds were also fixed at specific time-points for immunostaining of hepatic markers. PHHs stained with a human albumin antibody are shown to the right. Top row scale bar is 100 µm and bottom row scale bars are 5 mm.

### Hepatocyte functional assessments

Culture supernatants were assayed for albumin using a sandwich enzyme-linked immunosorbent assay (ELISA) kit with horseradish peroxidase detection (Bethyl Laboratories, Montgomery, TX) and 3,3’,5,5’-tetramethylbenzidine (TMB, Rockland Immunochemicals, Boyertown, PA) as the substrate. Absorbance values were quantified on the Synergy H1 multi-mode plate reader (BioTek, Winooski, VT). Cytochrome P450 2A6 (CYP2A6) enzyme activity was measured by incubating the cultures with 50 µM coumarin (Sigma-Aldrich) for 3 hours. The metabolite, 7-hydroxycoumarin (7-HC, Sigma-Aldrich), generated from coumarin was quantified via fluorescence detection (excitation/emission: 355/460 nm) using a standard curve on the Synergy H1 multi-mode plate reader.

### Cell staining

Scaffolds were fixed in 10% formalin solution (Sigma-Aldrich) overnight at 4°C and rinsed with 1X PBS. Fixed samples were embedded in paraffin following a series of xylene and graded ethanol incubations and cut into 8 μm thickness vertical cross-sections. Prior to cell staining, sections were deparaffinized. The fixed cultures were permeabilized and blocked using 0.3% Triton X-100 (Amresco) with 1% bovine serum albumin (BSA, Fisher Scientific) in 1X PBS for 4 hours at room temperature. Rabbit anti-human albumin (1:100) (Rockland Immunochemicals) or Mouse anti-human CYP3A4 (1:100) (GeneTex, Irvine, CA) antibodies were added with 0.3% Triton X-100 with 0.1% BSA in 1X PBS (dilution solution) and incubated overnight at 4°C. Cultures were then washed three times with 1X PBS. Cultures were labeled with either goat anti-rabbit IgG (H + L) Alexa Fluor 555 antibody (1:100) (Thermo Fisher Scientific) or rabbit anti-mouse FITC antibody (1:100) (GeneTex) in dilution solution for 3 hours at room temperature. After three 1X PBS washes, cultures were imaged using an Olympus IX83 microscope (Olympus America, Center Valley, PA).

### Data analysis

All findings were confirmed in 2-3 independent experiments (3-4 wells per condition and per experiment). Data processing was performed using Microsoft Excel. GraphPad Prism (La Jolla, CA) was used for displaying results. Mean and standard deviation are displayed for all data sets. Statistical significance was determined using Student’s t-test or one-way ANOVA followed by a Bonferroni pair-wise posthoc test (p< 0.05).

## RESULTS

### Porous, composite silk-collagen type I scaffolds support long-term PHH function

PHHs are an adherent cell type that require ECM proteins for optimal function; type I collagen extracted from rat tails (herein referred to as collagen I) is a popular ECM protein choice for PHH culture due to its abundant and cost-effective availability. Therefore, here we determined the optimal process of incorporating collagen I into our porous silk scaffolds. Porous silk scaffolds were initially prepared as described in methods, where silk fibroin solutions were frozen at −20°C, lyophilized at either −20°C or −80°C, and rendered insoluble by autoclaving to induce β-sheet formation. Silk was combined with collagen I to enable PHH attachment using three different methods. First, insoluble (autoclaved) silk scaffolds generated as above were incubated with a 1 mg/mL collagen I in 1X PBS solution for 2 hours at 37°C to enable passive collagen I adsorption onto the scaffolds (herein referred to as ‘silk + incubated collagen’). Second, lyophilized silk was autoclaved directly in a 1 mg/mL collagen I in 1X PBS solution (herein referred to as ‘silk + autoclaved collagen’). Third, 1 mg/mL collagen I in 1X PBS solution was added to the silk fibroin solution 3% (w/v) *prior* to its freezing and lyophilization; then, the silk scaffold containing incorporated collagen was further autoclaved in a 1 mg/mL collagen I in 1X PBS solution (herein referred to as ‘collagen incorporated silk + autoclaved collagen’). After fabrication, silk scaffolds were partially dehydrated by manually squeezing out the hepatocyte culture medium and the PHH suspension was added to the constructs (**Fig. 1**). Thereafter, PHH attachment and function were assessed in the various porous silk/collagen scaffolds.

The ‘silk + incubated collagen’ did not yield adequate PHH attachment nor function (data not shown) and thus was not used for further studies. When PHHs were seeded onto ‘silk + autoclaved collagen’ or ‘collagen incorporated silk + autoclaved collagen’ scaffolds, sufficient levels of PHH attachment were observed. To visualize PHH attachment in the scaffolds, the scaffolds were fixed, embedded in paraffin, and sectioned for staining over 1, 3, and 5 weeks of culture. PHHs within silk scaffolds stained positively for albumin (**Fig. 2** and **Supplemental Fig**. and CYP3A4 enzyme (**Supplemental Fig. 2**). While PHHs were found throughout the scaffolds, more cells were found closer to the exterior of the scaffolds with the small pores (∼73 µm +/- standard deviation of 25 µm) as can be expected for a passive delivery to cells to a porous scaffold. On the other, PHHs were more homogeneously distributed in the scaffolds with the large pores (235 µm +/- standard deviation of 84 µm). However, regardless of the pore size, PHHs remained stably housed within the ‘silk + autoclaved collagen’ or ‘collagen incorporated silk + autoclaved collagen’ scaffolds for several weeks.

**Figure 2.**
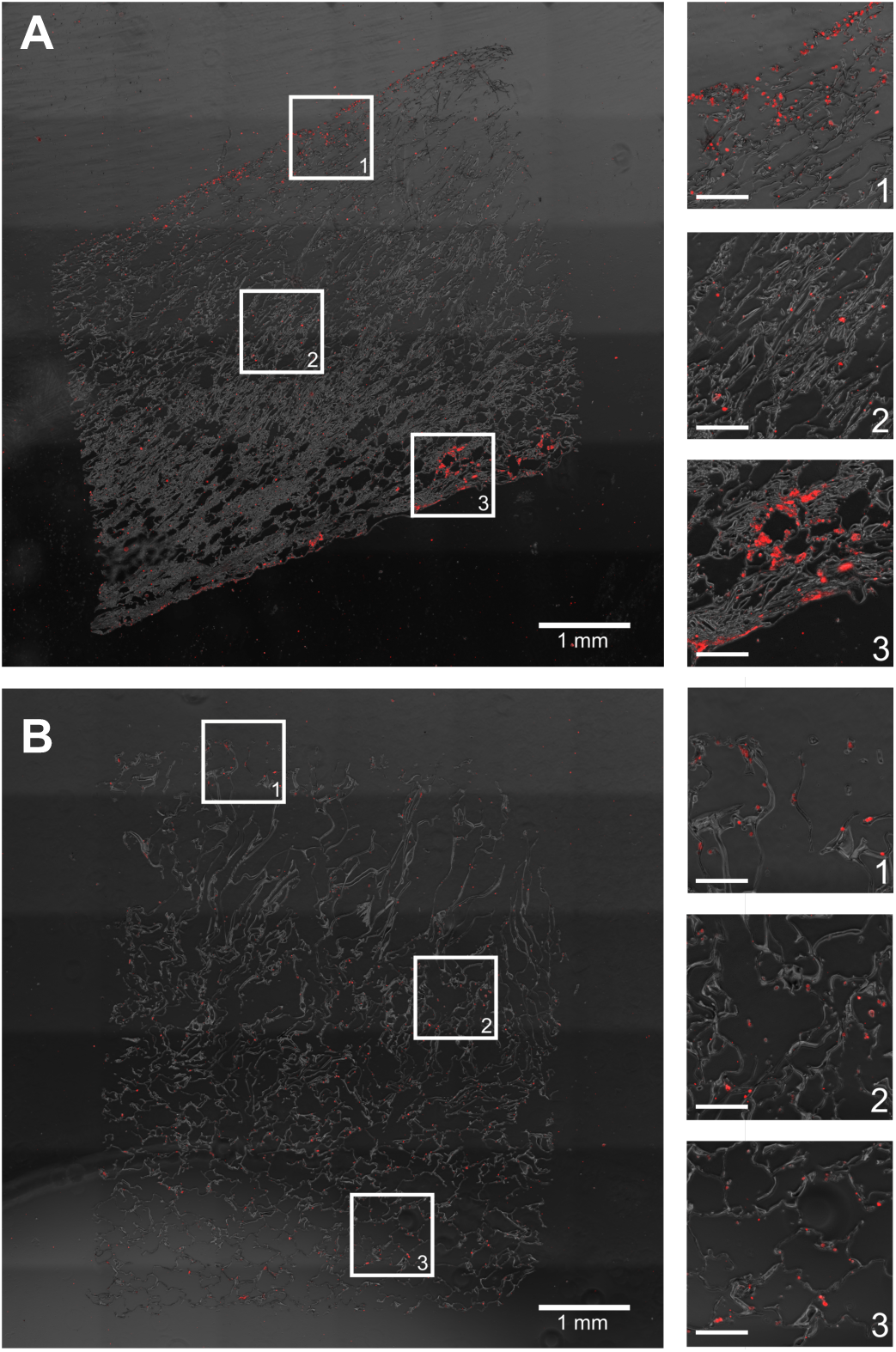
PHH distribution within composite silk-collagen I scaffolds of small or large pore sizes. Silk scaffolds containing collagen I were fabricated with (A) small pores (73 +/- 25 µm) or (B) large pores (235 +/- 84 µm) and then PHHs were seeded into the scaffolds per the protocols described in methods. After 3 weeks, the cell-laden scaffolds were fixed, sectioned, and stained for intracellular albumin. Shown in both panels is a single section and magnified images from three different regions within the scaffold to show the distribution of PHH attachment. The scale bars in magnified images are 250 µm.

At a functional level, PHHs within ‘collagen incorporated silk + autoclaved collagen’ scaffolds had ∼1.2- to 1.5-fold higher albumin production and ∼1.7- to 2.3-fold higher CYP2A6 enzyme activity than the ‘silk + autoclaved collagen’ scaffolds for ∼1 month in culture (**Fig. 3A**). Therefore, due to the consistent upregulation of PHH function over several weeks in ‘collagen incorporated silk + autoclaved collagen’ scaffolds, we selected this type of scaffold for all subsequent studies. With respect to the pore size, PHHs within ‘collagen incorporated silk + autoclaved collagen’ scaffolds with small pores displayed ∼1.5 to 1.8-fold higher albumin production and ∼2.2 to 3.8-fold higher CYP2A6 enzyme activity than the corresponding scaffolds with large pores for ∼1 month in culture (**Fig. 3B**). Finally, PHHs displayed ∼4- to 4.8-fold higher albumin production in 3D ‘collagen incorporated silk + autoclaved collagen’ scaffolds (small pore size) than when cultured as conventional monolayers on collagen-coated tissue culture plastic (2D control); additionally, CYP2A6 activity in PHHs was detected only in the 3D silk scaffolds and not in the conventional 2D control (**Supplemental Fig. 3A**).

**Figure 3.**
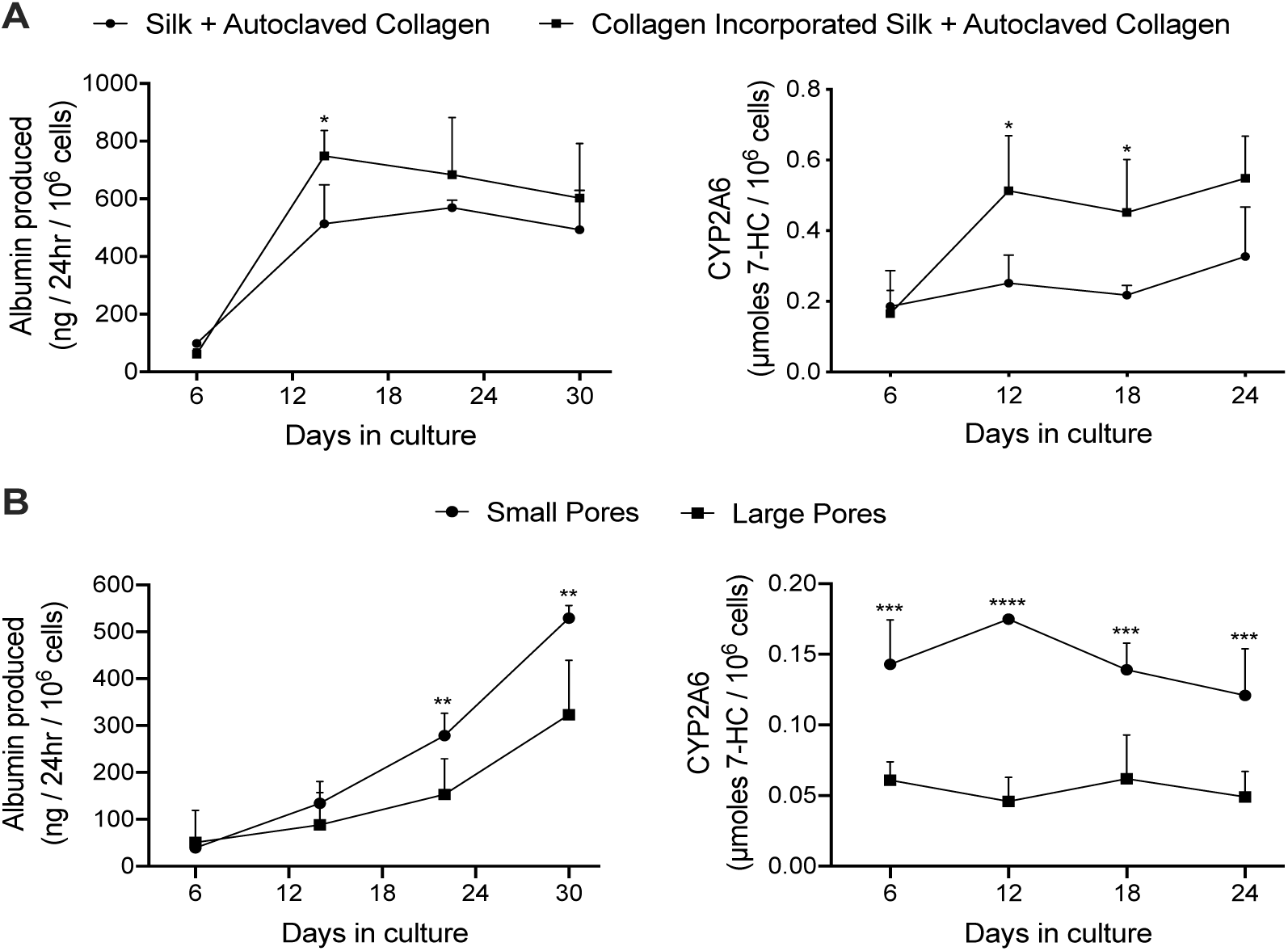
PHH functions within silk/collagen scaffolds of different pore sizes. Collagen I was incorporated into silk scaffolds by either autoclaving lyophilized silk with a 1 mg/mL collagen in PBS solution (‘Silk + Autoclaved Collagen’) or by mixing the solubilized silk solution with 1 mg/mL collagen in PBS followed by lyophilization and then further autoclaving in 1 mg/mL collagen in PBS (‘collagen incorporated silk + autoclaved collagen’). PHHs were seeded into the scaffolds as described in methods. (A) PHH albumin production and CYP2A6 enzyme activity over time in the two types of silk/collagen scaffolds with small pores (73 +/- 25 µm). (B) PHH albumin production and CYP2A6 activity over time in ‘collagen incorporated silk + autoclaved collagen’ scaffolds with either small or large pores (235 +/- 84 µm). \**p* ≤ 0.05, **p ≤ 0.01, ***p ≤ 0.001, and ****p ≤ 0.0001.

### Co-culture with fibroblasts enhances PHH functions within silk scaffolds for 5 months

3T3-J2 murine embryonic fibroblasts have been shown to induce functions in PHHs in both 2D and 3D (PEGDA-RGDS) culture formats [17, 29]. Therefore, here we sought to determine if a similar outcome could be achieved within the optimized porous silk/collagen scaffolds. The fibroblasts and PHHs were seeded at a 1:1 ratio into the ‘collagen incorporated silk + autoclaved collagen’ scaffolds (small pore size). In such co-cultures, PHHs remained within the scaffolds for at least 5 weeks and stained positive for both albumin (**Supplemental Fig. 1**) and CYP3A4 (**Supplemental Fig. 2**). The presence of the fibroblasts within the scaffolds enhanced PHH albumin production by ∼1.2 to 5-fold and CYP2A6 enzyme activity by ∼1.6 to 3.4-fold over ∼1 month of culture (**Fig. 4**).

**Figure 4.**
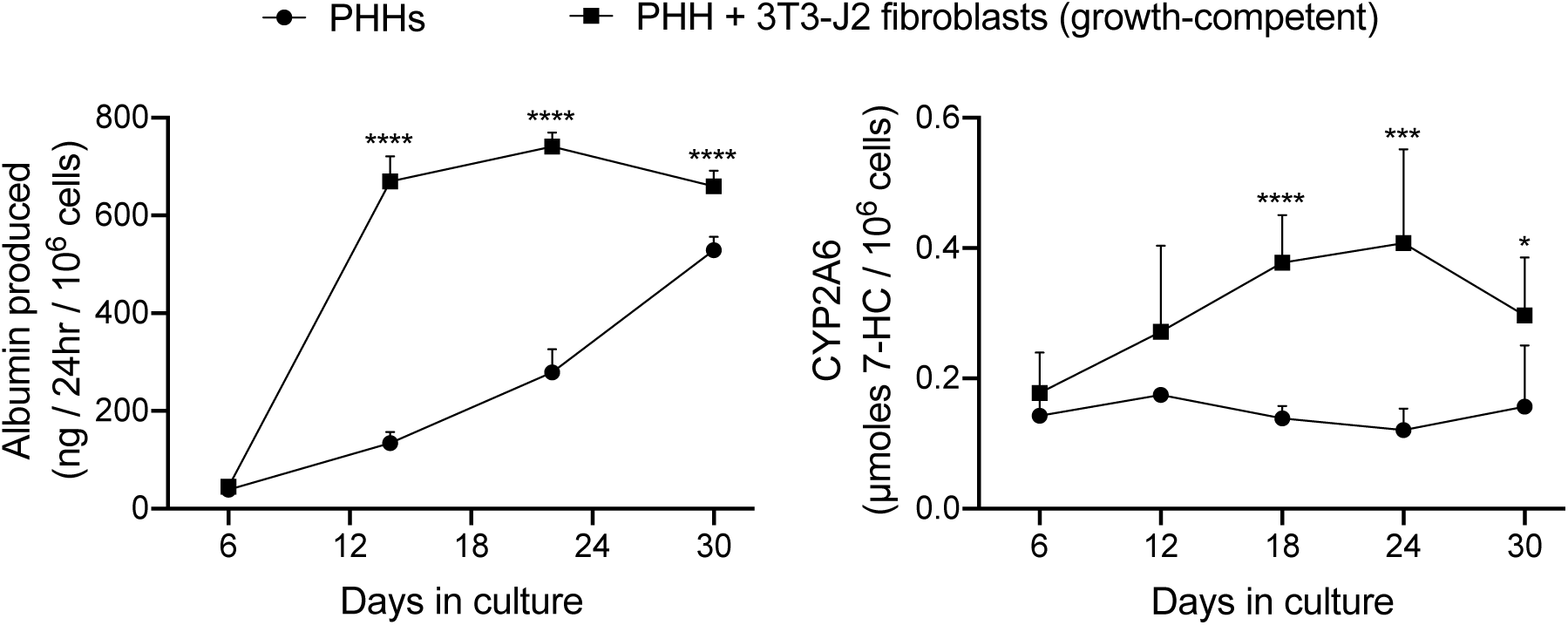
Functions of PHH-only and PHH + 3T3-J2 fibroblast co-cultures within silk/collagen scaffolds. ‘Collagen incorporated silk + autoclaved collagen’ scaffolds with small pores (73 +/- 25 µm) were fabricated as described in methods. A suspension of either PHH alone (500K cells per scaffold) or PHH + growth-competent 3T3-J2 fibroblasts (1:1 ratio, 500K cells for each type) was seeded into the scaffolds. Albumin production and CYP2A6 enzyme activity over time are shown. \**p* ≤ 0.05, ***p ≤ 0.001, and ****p ≤ 0.0001.

Due to concerns over fibroblast overgrowth within silk scaffolds in an implantation scenario in the future, the fibroblasts were growth-arrested via a preincubation with mitomycin-C prior to mixing with PHHs and seeding into the ‘collagen incorporated silk + autoclaved collagen’ scaffolds of either small or large pore sizes; PHH function was then tracked for up to 5 months in these co-cultures containing the growth-arrested fibroblasts while using the co-cultures containing the growth-competent fibroblasts as controls. As the growth-competent fibroblasts, the growth-arrested fibroblasts induced both albumin production and CYP2A6 activity in PHHs within both small (**Fig. 5A**) and large (**Fig. 5B**) pore scaffolds. While functions in both types of co-cultures were statistically similar within the large pore scaffolds, the growth-arrested fibroblasts induced ∼1.4 to 3.8-fold higher PHH albumin secretion in the small pore scaffolds after the first month of culture, whereas CYP2A6 activity was generally similar in both types of co-cultures across both types of pore sizes (**Fig. 5B**). Furthermore, both growth-competent and growth-arrested fibroblasts induced PHH functions maximally over ∼1 month of culture in both small pore or large pore size scaffolds and then the function declined over the next 15-30 days to 20-25% of the maximal levels, following which it declined more slowly over the remaining 2-3 moths of culture. Interestingly, co-cultures with both growth-competent and growth-arrested fibroblasts displayed more stable and higher (∼1.4- to 2.3-fold) albumin production in the large pore scaffolds in the fourth and fifth months of culture as compared to the small pore scaffolds; in contrast, CYP2A6 activity was similar in both types of co-cultures across both pore sizes for the entire duration of the culture time-period.

**Figure 5.**
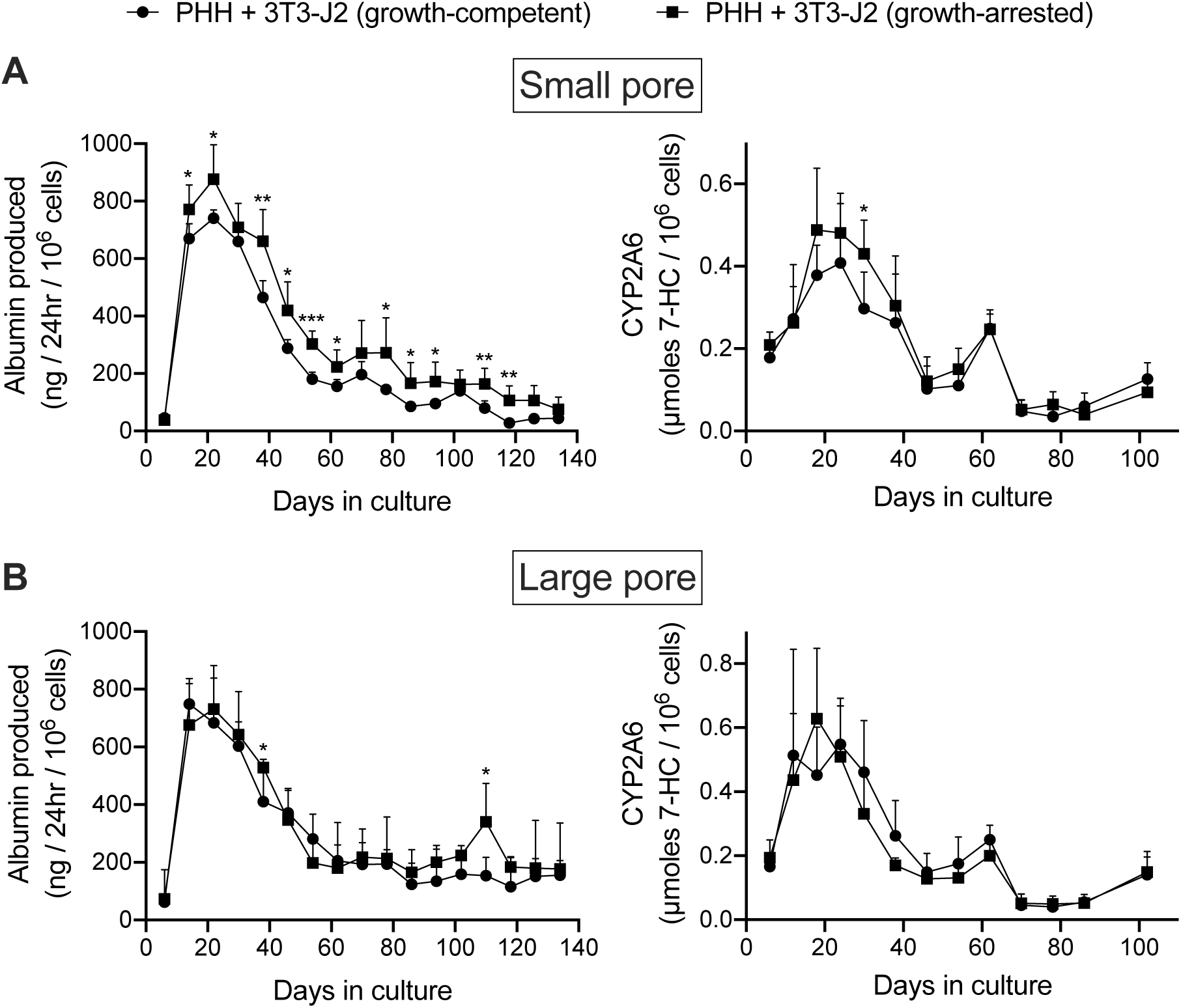
Long-term functions of PHH co-cultured with either growth-competent or growth-arrested 3T3-J2 fibroblasts in silk/collagen scaffolds of different pore sizes. ‘Collagen incorporated silk + autoclaved collagen’ scaffolds with either (A) small (73 +/- 25 µm) or (B) large (235 +/- 84 µm) pores were fabricated as described in methods. Albumin production and CYP2A6 activity are shown for almost 5 months for the different culture models. \**p* ≤ 0.05, **p ≤ 0.01, and ***p ≤ 0.001.

In contrast to co-cultures, PHHs alone displayed significantly lower (up to 5-fold lower for albumin and up to 10-fold lower for CYP2A6) levels of functions (**Supplemental Fig. 4**) than the co-cultures (**Fig. 5**); furthermore, while low levels of albumin was detected in the supernatants for 4 months, no CYP2A6 activity was detected after 2 months in the scaffolds with only PHHs, which was in contrast to the co-cultures.

When compared to conventional (2D) PHH + 3T3-J2 (growth-arrested) co-cultures on collagen-coated plastic, the co-cultures in the 3D ‘collagen incorporated silk + autoclaved collagen’ produced lower albumin in the first week of culture; however, albumin production in both models was similar from week 2 to week 6 of culture or ∼1.2-fold higher in the 3D scaffolds at certain time-points (**Supplemental Fig. 3B**). As with albumin, CYP2A6 activity in the 2D co-cultures was higher than the silk scaffolds for the first two weeks in culture; however, CYP2A6 activity in the 3D scaffolds became ∼1.3-and 2.1-fold higher between week 3 and 5 in culture than the 2D controls, and activity was only detected in 3D scaffolds and not in the 2D controls by week 6.

### Encapsulating co-cultures in Matrigel enhances PHH functions within silk scaffolds

The functions of PHH monolayers on adsorbed collagen have been shown to be enhanced when the cells are overlaid with gelled Matrigel™ [31]. Therefore, here we determined whether encapsulating PHH + 3T3-J2 (growth-arrested) co-cultures within growth factor reduced Matrigel (4 mg/mL) prior to dispensing into the silk scaffolds would further enhance PHH functions. Co-cultures were suspended in an ice-cold 4 mg/mL Matrigel (growth factor reduced) solution, which was then dispensed into the partially dehydrated ‘collagen incorporated silk + autoclaved collagen’ scaffolds of either small or large pore sizes (herein referred to as ‘Silk + Matrigel’). Control models included equal volumes of co-cultures/Matrigel suspensions dispensed directly into 48-well plates (referred to as ‘Matrigel’) or co-cultures suspended in an equal volume of culture medium (no Matrigel) dispensed directly into the above silk/collagen scaffolds (referred to as ‘Silk’).

As with the ‘Silk’ and ‘Matrigel’ control conditions, PHHs remained within the ‘Silk + Matrigel’ condition for at least 5 weeks and stained positive for both albumin (**Supplemental Fig. 1**) and CYP3A4 (**Supplemental Fig. 2**). Morphologically, co-cultures contracted the ‘Matrigel’ control scaffold whereas the conditions with silk in them did not show any appreciable contraction (not shown). At a functional level, co-cultures in the ‘Silk’ control condition with small pores displayed ∼1.2- to 2.2-fold higher albumin production and similar CYP2A6 enzyme activity than the ‘Matrigel’ control between week 2 and week 5 in culture, respectively (**Fig. 6A**). Furthermore, co-cultures in the ‘Silk + Matrigel’ condition with small pores displayed ∼1.1- to 1.6-fold and ∼1.5- to 2.6-fold higher albumin production for 5 weeks in culture than the ‘Silk’ and ‘Matrigel’ control conditions, respectively; similarly, co-cultures in the ‘Silk + Matrigel’ condition with small pores displayed ∼1.9- to 2-fold higher CYP2A6 activity between week 2 and week 5 in culture than the ‘Silk’ and ‘Matrigel’ control conditions (**Fig. 6A**).

**Fig. 6.**
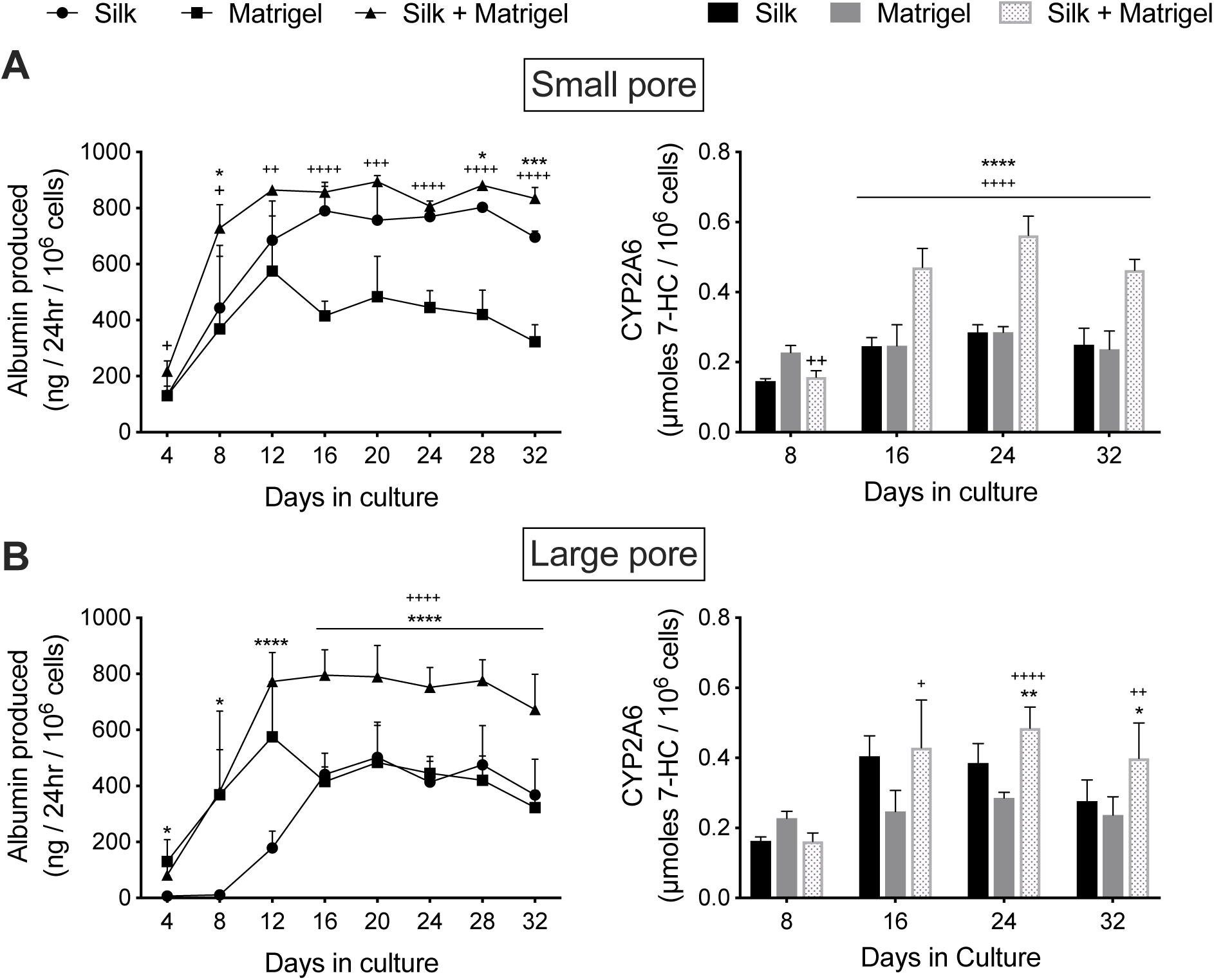
Functions of PHH + 3T3-J2 co-cultures in silk/collagen scaffolds, polymerized Matrigel only, and silk/collagen that contained polymerized Matrigel. ‘Collagen incorporated silk + autoclaved collagen’ scaffolds with (A) small (73 +/- 25 µm) pores or (B) large (235 +/- 84 µm) pores were fabricated as described in methods. The PHH + 3T3-J2 (growth-arrested, 1:1 ratio) co-cultures were either suspended in culture medium or a 4 mg/mL solution of cold Matrigel (growth factor reduced). Next, the co-cultures in culture medium or in Matrigel were dispensed into the silk scaffolds as described in methods and referred to as ‘Silk’ and ‘Silk + Matrigel’ conditions, respectively. Additionally, PHHs in Matrigel was dispensed directly into the wells of a 48-well plate as a control condition (‘Matrigel’). Upon incubation at 37°C, the Matrigel polymerized, thereby encapsulating the co-cultures. Albumin production and CYP2A6 activity are shown over time for the different culture models. \**p* ≤ 0.05, **p ≤ 0.01, ***p ≤ 0.001, and ****p ≤ 0.0001 for ‘Silk +Matrigel’ vs. ‘Silk’. ^+^*p* ≤ 0.05, ^++^p ≤ 0.01, ^+++^p ≤ 0.001, and ^++++^p ≤ 0.0001 for ‘Silk +Matrigel’ vs. ‘Matrigel’.

Co-cultures in the large pore ‘Silk’ control initially showed for the first 2 weeks lower albumin production and lower CYP2A6 activity than the ‘Matrigel’ control; however, between week 3 and week 5, co-cultures in the ‘Silk’ condition produced relatively similar levels of albumin and displayed ∼1.2- to 1.6-fold higher CYP2A6 activity than the ‘Matrigel’ control (**Fig. 6B**). Furthermore, co-cultures in the ‘Silk + Matrigel’ condition with large pores displayed ∼1.6- to 4.3-fold and ∼1.3- to 2.1-fold higher albumin production between week 2 and week 5 in culture than the ‘Silk’ and ‘Matrigel’ control conditions, respectively; similarly, co-cultures in the ‘Silk + Matrigel’ condition with large pores displayed ∼1.1- to 1.4-fold and ∼1.7-fold higher CYP2A6 activity between week 2 and week 5 in culture than the ‘Silk’ and ‘Matrigel’ control conditions, respectively (**Fig. 6B**).

Finally, when a type I collagen gel (4 mg/mL) was used instead of Matrigel to embed PHH + 3T3-J2 (growth-arrested) co-cultures prior to dispensing into the ‘collagen incorporated silk + autoclaved collagen’ scaffolds (small pores), unlike the ‘Matrigel’ control, the ‘Collagen’ control was not contracted to any appreciable degree, which was similar to the ‘Silk’ control and ‘Silk + Collagen’ condition (not shown). At a functional level, co-cultures in the ‘Collagen’ control displayed statistically similar albumin production and CYP2A6 activity as the ‘Silk’ condition and ‘Silk + Collagen’ condition over 4-5 weeks in culture (**Supplemental Fig. 5**).

## DISCUSSION

While some biomaterials have been explored to fabricate implantable human liver tissues towards addressing the critical shortage of donor organs for transplantation, there remains a need to further explore PHH interactions with different biomaterials towards determining optimal conditions for their long-term survival *in vitro* and eventual translation to humans. Scaffolds created using silk proteins have highly desirable properties (i.e. biocompatibility, tunable mechanical properties, degradation rates, and low immunogenicity) for fabricating implantable tissues. Here, we show for the first time that porous composite silk-collagen I scaffolds can be used to sustain PHH functions for 5 months *in vitro* in the presence of supportive fibroblasts.

Silk proteins have been molded into a variety of shapes, such as hydrogels, tubes, films, and sponges [32]. Here, we selected porous silk (fibroin) sponges since they allow culture medium to penetrate into the interior of the scaffolds to maintain cell viability even without perfusion. Furthermore, since silk proteins do not support rapid cell attachment, we combined the silk with rat tail collagen I, which is widely used to enable PHH attachment to various substrates due to its availability in large quantities and cost-effectiveness relative to human-derived proteins [33]. Collagen from different species is well tolerated *in vivo* [34]; however, it has weak mechanical properties and is rapidly degraded post-implantation [35]. Therefore, combining collagen I with mechanically robust silk that can be tuned for degradation rates is a proven strategy [22].

Collagen I was incorporated into the silk scaffolds using three different methods. First, insoluble scaffolds were incubated with a solution of collagen I, similar to the strategy used for tissue culture plastic [33]; however, PHHs did not attach well, suggesting that collagen did not adsorb at adequate levels onto the silk scaffolds. In contrast, collagen I incorporated into the silk during its β-sheet (crystal) formation induced by autoclaving led to robust PHH attachment for several weeks as verified by immunostaining for PHH markers, albumin and CYP3A4; such an effect may be due to integration of the collagen I into the β-sheets. Further adding collagen during the lyophilization of the solubilized silk solution and then also autoclaving in the collagen I solution similarly promoted PHH attachment. While PHHs attached within silk/collagen scaffolds with both small (73 +/- 25 µm) and large (235 +/- 84 µm) pores, the distribution of cell attachment was more uniform in the latter as can be expected from a passive (versus via a bioreactor) cell seeding process used here. Nonetheless, scaffolds with both pore sizes retained PHHs for several weeks as verified by immunostaining.

To appraise PHH functionality, we measured albumin production, a marker of liver’s synthetic capability, and CYP2A6 enzyme activity as a marker of liver’s drug metabolism capacity. While CYP3A4 is the most abundant CYP450 enzyme in the liver, we were unable to utilize a well-established luminescent assay [29] to assess this enzyme due to a lack of assay performance, which may be due to the binding of the luminescent substrate to the silk proteins; however, PHHs stained for CYP3A4 protein for several weeks in the silk/collagen scaffolds. In contrast, the conversion of coumarin by CYP2A6 into 7-hydroxycoumarin could be readily assessed within silk scaffolds, which tracks similarly to CYP3A4 activity in PHH cultures as we have shown previously [29]. Overall, PHHs cultured within the optimal ‘collagen incorporated silk + autoclaved collagen’ scaffolds had up to 2.3-fold higher function than the ‘silk + autoclaved collagen’ scaffolds, suggesting that the addition of collagen during lyophilization and autoclaving is beneficial for PHHs; however, it is not clear if such an effect is due to better retention of PHHs over time within the optimal scaffolds or higher functions on a per cell basis, which we plan to elucidate in future studies. Nonetheless, PHHs in the optimal silk/collagen scaffolds retained functions for at least 1 month and were used for all subsequent studies.

The optimal silk/collagen scaffolds with small pore sizes induced up to ∼4-fold higher function than scaffolds with larger pores; such an effect could be due to better promotion of PHH homotypic interactions within the smaller pores as was observed with staining. Indeed, the promotion of homotypic interactions is critical for stabilizing PHH phenotype in both 2D (e.g. confluent monolayers) and 3D (e.g. spheroids) configurations previously [33]. Finally, CYP2A6 activity was not detectable in 2D monocultures on collagen-coated plastic but was well-retained in the 3D silk/collagen scaffolds for several weeks, which is likely due to the promotion of homotypic interactions and spheroidal morphology of PHHs with minimal spreading since hepatocyte spreading inversely correlates with differentiated functions [36].

Co-culture with liver- and non-liver-derived non-parenchymal cell (NPC) types can positively regulate the survival and functions of hepatocytes in both the pre- and postnatal livers [33, 37]. Such interactions have been long replicated *in vitro* by co-culturing primary hepatocytes with various liver- and non-liver-derived NPCs, including C3H/10T1/2 mouse embryo cells [38], 3T3-J2 murine embryonic fibroblasts [39], and human fibroblasts [40]. The 3T3-J2 fibroblasts, in particular, have been shown to express key liver-like molecules, such as decorin [41] and T-cadherin [42], which help these fibroblasts induce higher levels of functions in PHHs in 2D co-cultures than primary human dermal fibroblasts (not shown), human liver-derived hepatic stellate cells [43], liver sinusoidal endothelial cells [44], and Kupffer cells [45]. The positive effects of 3T3-J2 on PHH functions have also been shown in 3D PEGDA-RGDS hydrogels [17]. However, it is not clear if such hydrogels can maintain the survival and function of PHH-3T3-J2 co-cultures *in vitro* beyond 1-2 weeks, which may be mitigated by the use of a silk/collagen scaffold that, unlike PEGDA-RGDS, allows cells to modulate their contacts and aggregate size over time and facilitates ECM deposition/remodeling due to the porous structure.

The presence of the 3T3-J2 fibroblasts within the optimal silk/collagen scaffolds enhanced PHH functions up to 5-fold relative to PHH-only controls. However, since the use of growth-competent fibroblasts may lead to fibrosis upon implantation, we also growth-arrested the fibroblasts using mitomycin-C (alkylates DNA) before co-culture with PHHs and showed that such a strategy does not affect the ability of the fibroblasts to induce PHH functions. Importantly, in contrast to PHH-only cultures, the effects of the fibroblasts on PHH functions in the optimal silk/collagen scaffolds lasted 5 months, which bodes well for the sustained shelf-life of such constructs for future on-demand implantation into animal models and eventually humans. However, co-culture functions in the scaffolds declined after the first month in culture, which suggests that further improvements are needed to the ECM composition within the scaffold and/or media formulation to maintain steady functional levels. Nonetheless, the longevity of PHH/3T3-J2 co-culture functions within silk/collagen scaffolds is unprecedented in this field.

When compared to conventional 2D PHH/3T3-J2 co-cultures on collagen-coated tissue culture plastic, co-cultures within the 3D silk/collagen scaffolds displayed lower functions in the first 1-2 weeks of culture; such as an outcome is likely due to the prolonged time it takes for 3D aggregates/spheroids of PHHs to adequately form and make sufficient cell-cell adhesions, as also observed in other spheroid generation platforms [46]. However, once the 3D co-cultures had adequately established within the optimal silk/collagen scaffolds, functions in 3D were up to 2-fold higher than 2D; importantly, CYP2A6 activity was detected only in 3D and not in 2D after 6 weeks in culture.

While PHH/3T3-J2 co-cultures housed within optimal silk/collagen scaffolds function for 5 months, the liver has many other ECM proteins that can modulate hepatic functions [47]; furthermore, hepatic functions are enhanced when cultured on or within soft hydrogels [33, 48]. Therefore, here we encapsulated PHH/3T3-J2 co-cultures in Matrigel, an ECM routinely used to modulate PHH functions in different culture formats [33]. The encapsulation was done either inside silk/collagen scaffolds or in plastic control wells. Morphologically, co-cultures within Matrigel alone contracted over time and detached from plate surfaces, which is a common problem with cell culture in soft gels. However, when Matrigel was injected into the silk/collagen scaffolds, the construct remained intact in size/shape and PHH functions ramped up faster and were up to 2.6-fold higher than the silk/collagen and Matrigel-only control conditions. In contrast, no synergistic increase in coculture function was observed when collagen I gels were entrapped within the silk/collagen scaffolds, suggesting that collagen I alone is not sufficient to promote the highest PHH functions. Therefore, combining silk scaffolds with ECM hydrogels, especially those with a liver-inspired and complex protein composition such as Matrigel or decellularized liver matrix, may ultimately be the most optimal strategy to create an implantable engineered human liver construct that leverages the advantages of both silk scaffolds and ECM hydrogels.

Previous investigations using silk scaffolds using either gelatin or RGD as integrin binding sites for culture of human liver cell lines (i.e., HepG2 cell line) or rodent hepatocytes have shown minimal inflammation and no adaptive immune response or fibrosis upon implantation into rodents models [23, 26]. While it is difficult to fully extrapolate these studies to our work due to the use of different cell types, these previous studies nonetheless show that implanting liver constructs made of silk/ECM scaffolds has promise as a potential therapy that needs further exploration.

While our studies culturing PHH and supportive fibroblasts for 5 months in silk/collagen scaffolds are unprecedented, further work needs to be done to create a clinically viable human liver tissue. First, incorporation of liver sinusoidal endothelial cells into specific vascular compartments within the silk scaffolds is likely necessary to allow better integration of scaffold vasculature with the host vasculature; indeed, hollow channels can be created by casting the silk scaffold around a mold made of polydimethylsiloxane and Teflon-coated wires as was done for intestinal cultures [27]. Inclusion of mesenchymal stem cells with endothelial cells in the silk scaffolds may allow better formation of stabilized neovessels as was done in Matrigel [49]. Second, while PHHs are most physiologically relevant and strategies are being developed to harness their tremendous expansion potential as *in vivo* using small molecule modulators of key pathways [9], their use will still necessitate immune-suppression in eventual recipients. While immune suppression via drugs is routinely practiced in organ transplantation, the use of iPSC-derived human liver cells will allow a nearly infinite source of autogenic cells. However, protocols to improve the differentiation of the iPSC-derived human liver cells, including hepatocyte-like cells, need to be significantly improved to induce adult-like functional levels before these cells can be used for routine cell therapy [8]. Finally, the liver has Kupffer cells (resident macrophages), hepatic stellate cells (vitamin A storing cells), and cholangiocytes (epithelial cells lining the bile ducts) that would need to be included in the scaffolds alongside PHHs and supportive fibroblasts to ensure a fully functioning liver for transplantation. Nonetheless, since PHHs perform the majority of liver’s functions, starting with their long-term culture within silk/collagen scaffolds is an important first step in the development of an implantable engineered liver construct, as demonstrated here.

In conclusion, silk/collagen porous scaffolds can maintain PHH functions for 5 months *in vitro* in the presence of growth-arrested 3T3-J2 fibroblasts; further encapsulating such co-cultures in Matrigel while housed within the silk/collagen scaffolds stabilizes the hydrogel and enhances PHH functions relative to silk/collagen or Matrigel-only controls. Ultimately, PHH/3T3-J2 co-cultures within silk/collagen scaffolds can serve as building blocks for strategies in regenerative medicine and to screen the efficacy and/or toxicity of lead compounds during drug development.

## AUTHOR CONTRIBUTIONS

D.A.K., W.L.S., D.L.K., and S.R.K. designed experiments; D.A.K. and W.L.S. executed experiments; D.A.K. and W.L.S. analyzed data. D.A.K., W.L.S., D.L.K., and S.R.K. wrote the manuscript.

## ACKNOWLEDGEMENTS

Funding for this work was provided by the National Science Foundation CAREER award (CBET-1351909) to S.R.K. W.L.S. acknowledges support from the National Institutes of Health (NIH) Institutional Research and Academic Career Development Awards program at Tufts University (K12GM074869, Training in Education and Critical Research Skills (TEACRS)) and we acknowledge support from NIH P41 (EB002520) for this work.

