## Supplemental Figure for "Primary Human Hepatocytes Maintain Long-term Functions in Porous Silk Scaffolds Containing Extracellular Matrix Proteins"

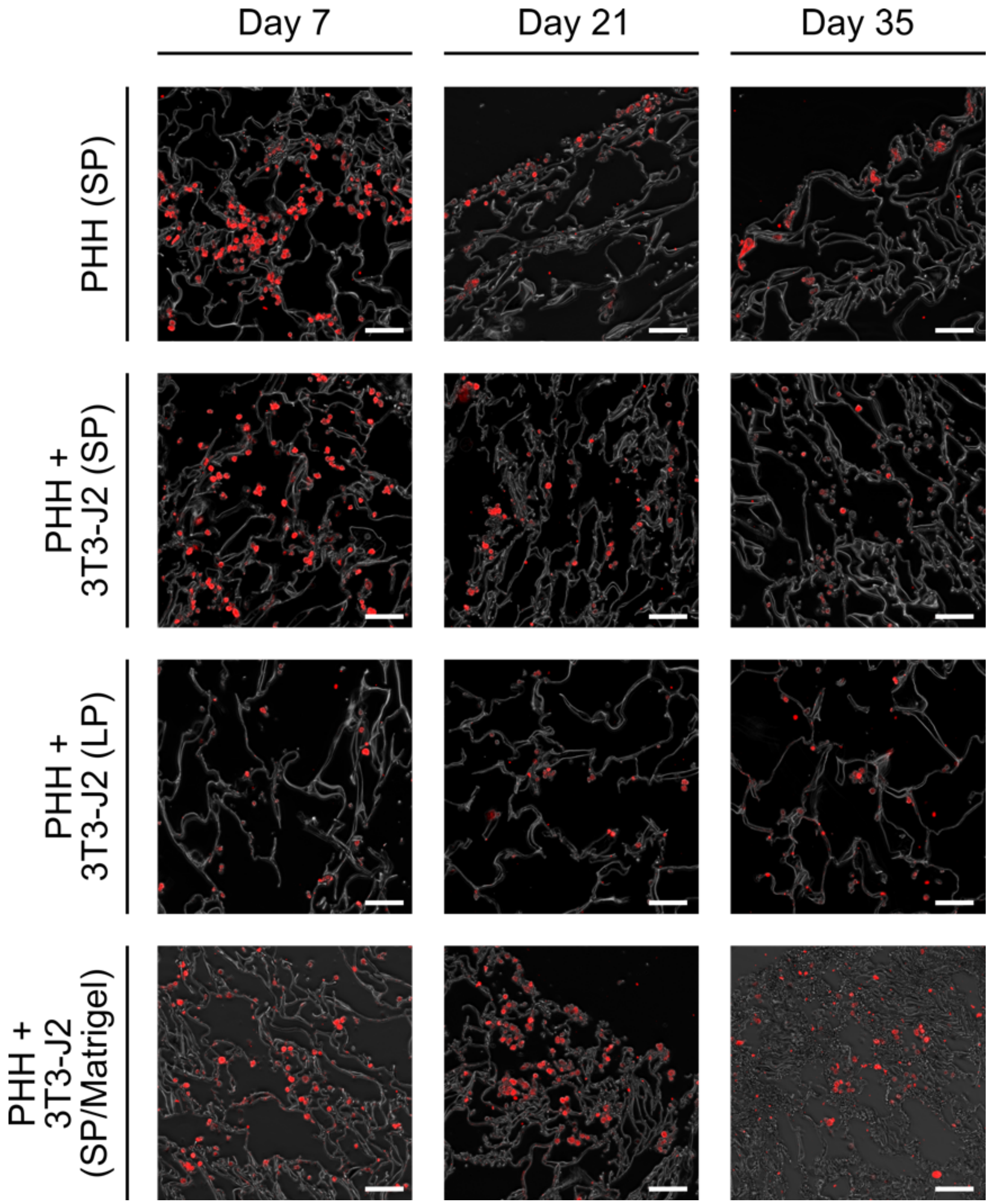


**Supplemental Fig. 1. PHH intracellular albumin stain within silk/collagen scaffolds.** PHH-only cultures and PHH + 3T3-J2 fibroblast co-cultures were seeded into 'collagen incorporated silk + autoclaved collagen' scaffolds of different pore sizes as described in methods of the main manuscript and cultured for 5 weeks *in vitro*. Conditions include PHH-only cultures in small pore silk [PHH(SP)], PHH + 3T3-J2 co-cultures in small pore silk [PHH + 3T3-J2(SP)] and large pore silk [PHH + 3T3-J2(LP)], and PHH + 3T3-J2 co-cultures in small pore silk with incorporated Matrigel hydrogel [PHH + 3T3-J2(SP/Matrigel)]. PHHs were positive for albumin (red) over the 5-week culture period. All scale bars are 100 µm.

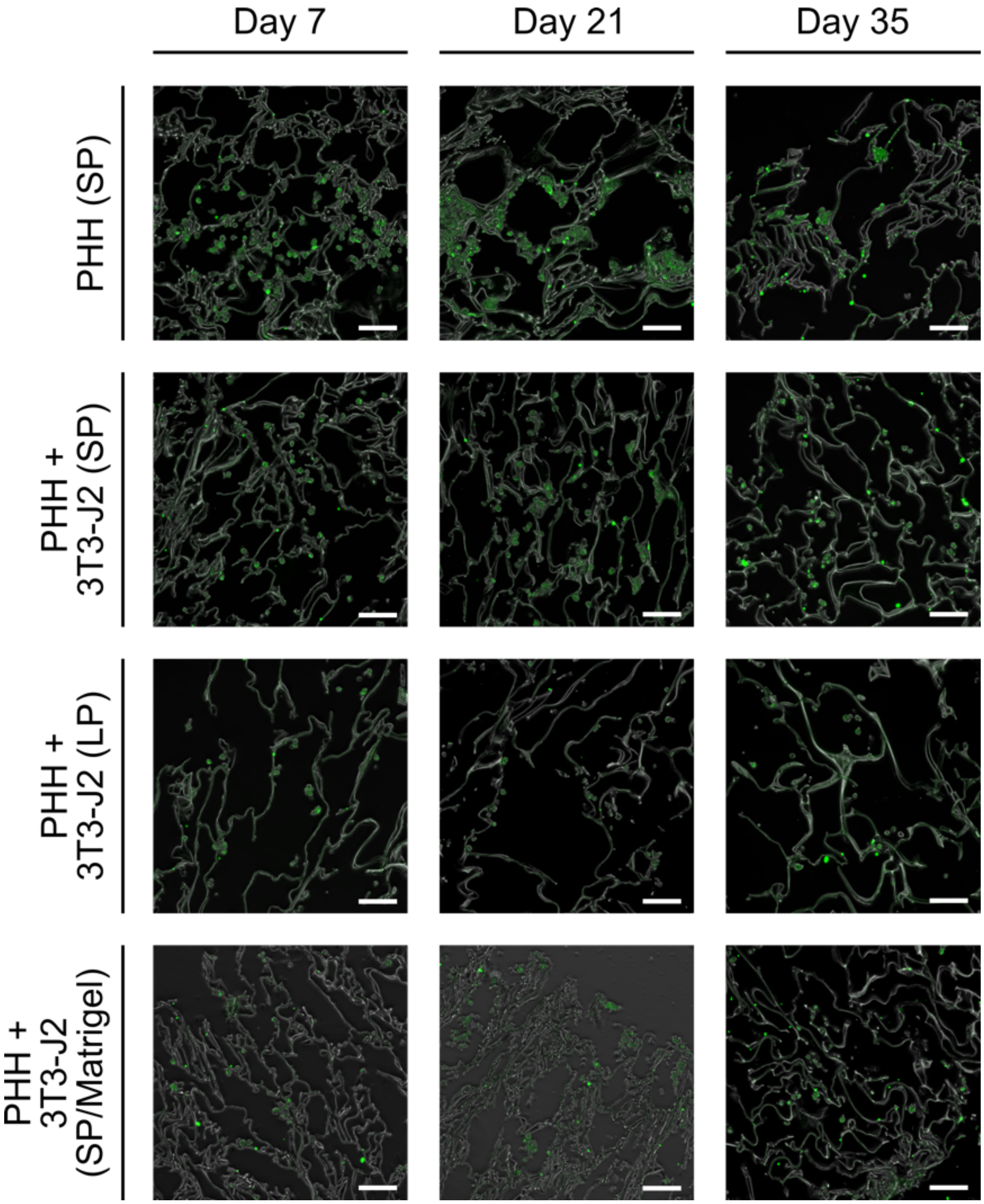


**Supplemental Fig. 2. PHH intracellular cytochrome P450 3A4 (CYP3A4) stain in silk/collagen scaffolds.** PHH-only cultures and PHH + 3T3-J2 fibroblast co-cultures were seeded into ‘collagen incorporated silk + autoclaved collagen’ scaffolds of different pore sizes as described in methods of the main manuscript and cultured for 5 weeks *in vitro*. Conditions include PHH-only cultures in small pore silk [PHH(SP)], PHH + 3T3-J2 co-cultures in small pore silk [PHH + 3T3-J2(SP)] and large pore silk [PHH + 3T3-J2(LP)], and PHH + 3T3-J2 co-cultures in small pore silk with incorporated Matrigel hydrogel [PHH + 3T3-J2(SP/Matrigel)]. PHHs were positive for CYP3A4 (green) over the 5-week culture period. All scale bars are 100 µm.

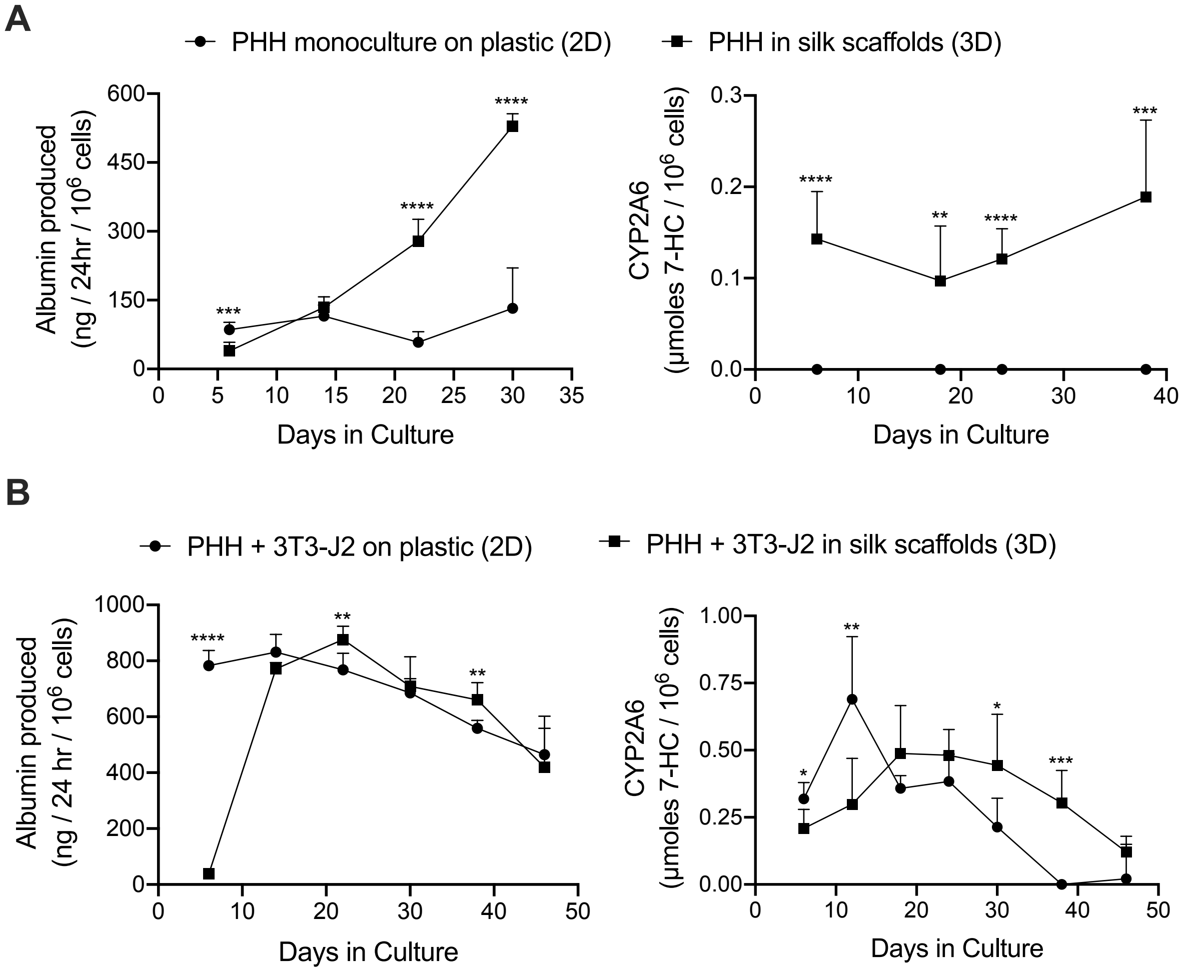


**Supplemental figure 3. PHH-only and PHH + 3T3-J2 functions in silk/collagen scaffolds (3D) and on collagen adsorbed tissue culture plastic (2D).** 'Collagen incorporated silk + autoclaved collagen' scaffolds with small (73 +/- 25 µm) pores were fabricated or collagen I was adsorbed onto tissue culture plastic wells as described in methods of the main manuscript. (A) PHH-only (monoculture) or (B) PHH + 3T3-J2 (growth-arrested, 1:1 ratio between the cell types) suspensions were seeded onto the 3D or 2D platforms at the same cell numbers and media volumes to enable comparisons. Albumin production and CYP2A6 activity are shown over time for the different culture models. *$p \leq 0.05$, **$p \leq 0.01$, ***$p \leq 0.001$, and ****$p \leq 0.0001$.

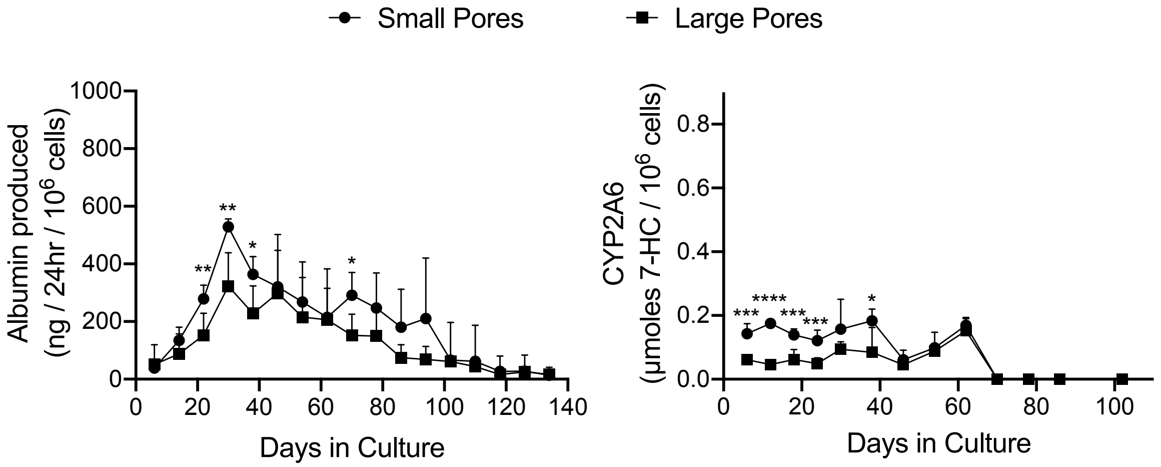


**Supplemental Figure 4. Long-term functions of PHHs in silk/collagen scaffolds of different pore sizes.** 'Collagen incorporated silk + autoclaved collagen' scaffolds with either (A) small (73 +/- 25 µm) or (B) large (235 +/- 84 µm) pores were fabricated as described in methods of the main manuscript. Albumin production and CYP2A6 activity are shown over time for PHHs within the scaffolds of two different pore sizes. **p* ≤ 0.05, **p ≤ 0.01, ***p ≤ 0.001, and ****p ≤ 0.0001.

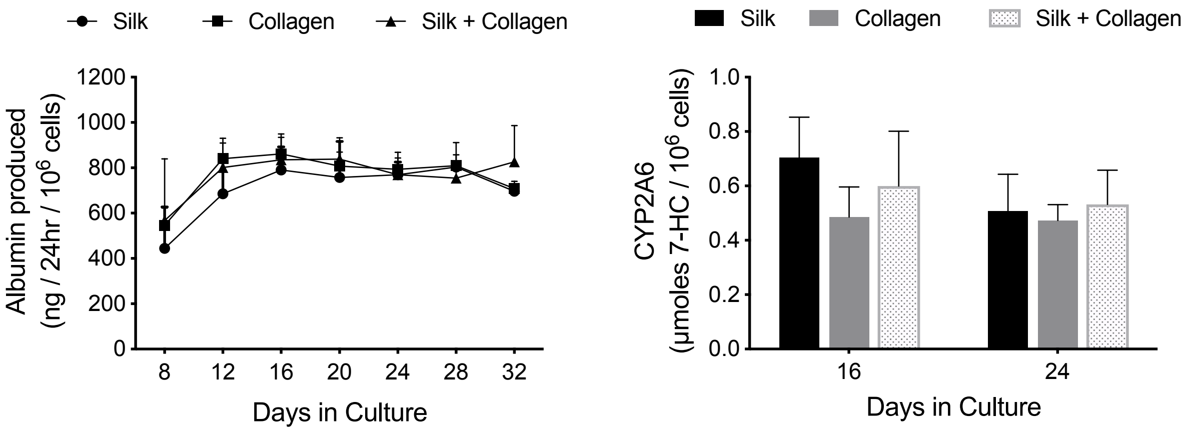


**Supplemental figure 5. Functions of PHH + 3T3-J2 co-cultures in silk/collagen scaffolds, polymerized collagen I, and silk/collagen + polymerized collagen I.** 'Collagen incorporated silk + autoclaved collagen' scaffolds with small (73 +/- 25 µm) pores were fabricated as described in methods of the main manuscript. The PHH + 3T3-J2 (growth-arrested, 1:1 ratio) co-cultures were either suspended in culture medium or a 4 mg/mL solution of cold rat tail collagen I. Next, the co-cultures in culture medium or in collagen were dispensed into the silk scaffolds as described in methods and referred to as 'Silk' and 'Silk + Collagen' conditions, respectively. Additionally, PHHs in collagen was dispensed directly into the wells of a 48-well plate ('Collagen' condition). Upon incubation at 37°C, the collagen polymerized, thereby encapsulating the co-cultures. Albumin production and CYP2A6 activity are shown over time for the different culture models.
